# Bioactive compounds from *Chrysosporium multifidum*, a fungus isolated from *Hermetia illucens* gut microbiota

**DOI:** 10.1101/669515

**Authors:** Yesenia Correa, Billy Cabanillas, Valérie Jullian, Daniela Álvarez, Denis Castillo, Cédric Dufloer, Beatriz Bustamante, Elisa Roncal, Edgar Neyra, Patricia Sheen, Michel Sauvain

**Author notes:** Corresponding autor.

## Abstract

The gut microbiota of insects contains a wide range of organisms that protect them against the attack of pathogens by releasing various types of bioactive compounds. In the present study, we report the isolation and identification of the fungus *Chrysosporium multifidum* as a component of the microbiota from the larval gut of *Hermetia illucens.* Extract from the broth culture of *C. multifidum* showed moderate activity on a strain of methicillin-resistant *Staphylococcus aureus* (MRSA). The bioguided isolation of the extract resulted in the characterization of six α-pyrone derivatives (**1**-**6**) and one diketopiperazines (**7**), among them 5,6-dihydro-4-methoxy-6-(1-oxopentyl)-2H-pyran-2-one (**4**) showed the best activity (IC_50_ = 11.4 ± 0.7 µg/ml and MIC = 62.5 μg/ml).

## Introduction

*Hermetia illucens*, also known as the black soldier fly (BSF), is native species of Americas. In the last decade, interest in this insect has increased because its larvae can reduce various types of organic waste [1]. BSF larvae have the ability to extract efficiently the protein and lipid content of the wastes they feed on. The larvae and their derivatives appear as a promising alternative in sustainable production for the animal feed industry and biodiesel production. Several research groups and companies around the world are conducting studies in order to optimize their production on a large scale [2]. BSF larvae can feed on different types of organic waste unaffected by bacteria entering their gut [3, 4]. The control of larvae over these pathogens would take place through the release of bioactive substances released into the larva [5]. Thus, compounds generated by BSF larvae could be an alternative to drugs used to treat bacteria harmful to humans, especially those that have shown increased antibiotic resistance in recent years [6, 7]. The gut microbiota arises as an important source of antibacterial compounds due to the existing evidence on the different biological activities found in the fungi associated with insects [8-13]. Recently, a study showed the diversity of fungi isolated from the gut of BSF larvae. Among them, *Trichosporon asahii* showed to be active on sensitive strains of *Candida glabrata* and *Candida lusitaniae* [14]. Other antibacterial substances such as peptides [15, 16] and lipids [17] have also been isolated from BSF, but information is still limited. The purpose of this work is to describe the isolation and identification of a fungus with antimicrobial activity from the gut of BSF larvae, as well as the identification of isolated compounds from the *in vitro* culture of the fungus.

## Material and methods

### Larvae rearing

The BSF larvae used in this experiment were obtained from the breeding colony established at the Universidad Peruana Cayetano Heredia (Lima, Peru) maintained at 28 ± 1°C and 70% of relative humidity. Specimens were fed with fresh unsterilized chicken guano for 11 days. After this time larvae exhibited lengths between 1.5 - 2 cm.

### Extraction of gut from larvae and isolation of active fungi

Samples were collected in triplicate; each collection corresponded to larvae obtained from a different breeding cycle. The larvae were washed with EtOH 70% and sterilized during 15 min with UV light. Each larva was dissected transversally and the gut was extracted. A sample of 0.5 cm was taken from the middle gut to isolate the fungi. The gut samples were homogenized in 200 µl of saline solution 0.89%. The resulting solutions were diluted (10^-1^ - 10^-4^) and 10 µl were seeded in plates containing potato dextrose agar (PDA) (BD-Difco®) and Sabouraud agar (SBA) (BD-Difco®) both supplemented with chloramphenicol (100 mg/L) and gentamicin (50 mg/). To confirm the absence of external contamination two controls were used, a first control using the saline solution used in homogenization, and a second control resulting from the swabbing of the larvae surface after disinfection and before dissection. Plates with sample dilutions were incubated for 21 days at 30 ± 1°C. Strains with different morphology were separated repeatedly and constantly in the same media to obtain pure colonies. The activity of fungi colonies to *Staphylococcus aureus* subsp. *aureus* ATCC 43300 and *Salmonella enterica* subsp. *enterica* var. Typhimurium ATCC 13311 were determinate by a previously described method [18], with small modifications. From all the 25 isolates obtained, one of them showed the best activity being submitted to DNA analysis to establish its identity at the specie level.

### DNA extraction from the active fungus

A sample of 100 mg of mycelium was transferred into a 2 ml tube and 700 µl of extraction buffer consisting of 0.1 M Tris-HCl (pH=8), 20 mM EDTA (pH=8), 1.4 M NaCl, 0.2% (v/v) 2-mercaptoethanol and 2% (w/v) CTAB was added with mixed acid-washed 150-212 µm glass beads. The mixture was placed inside a Quiagen Tissue Lyser II for 30 seconds, and aliquot of 15 µl RNAsa A (20 mg/ml) was added and then mixed at 55°C and 850 rpm for 30 min. After, 700 µl of chloroform:isoamyl alcohol (24:1) was added, and the mixture was centrifuged at 14,000 rpm for 10 min at room temperature. The supernatant was mixed with 50 µl of 10% (w/v) CTAB and 600 µl of chloroform and then centrifuged at 14,000 rpm for 10 min. The new supernatant was transferred to a clean 1.5 ml tube an equal volume of ice-cold was added, leaved at −20°C for one night, and then centrifuged at 14,000 rpm for 20 min. The pellet was washed with 1 ml of 80% ethanol (4°C) and centrifuged at 14,000 rpm for 10 min, the operation was repeated and the resulting pellet was left to dry at room temperature for 3 hours. The quantification of DNA was carried out by Nanodrop. Finally, an electrophoresis was performed to verify the integrity of the DNA in 1.5% agarose gel. The extracted DNA was stored at −20°C until use.

### PCR amplification and sequencing of the active fungus DNA

A fungal DNA amplification was performed by conventional PCR, two zones were chosen for the amplification, the first one amplified an area covering ITS1-2 rRNA with the universal primers ITS1 (TCCGTAGGTGAACCTGCGGG) and ITS4 (TCCTCCGCTTATTGATGGC); while the second covered D1/D2 domains of large subunit (LSU) ribosomal DNA(rDNA) with the universal primers NL1 (GCATCAATAAGCGGGAGGAAAG) and NL4 (GGTCCGTGTTTCAAGGGG. A master mix solution was prepared containing 25 µl of KOD (Hot start Master Mix-Sigma Aldrich), 1.5 µl of each primer and 18.5 µl of NFW. The DNA was included in 3.5 µl at a concentration of 20 ng/µl. Cycling conditions were as follow: initial denaturation at 94°C for 5 min, followed by 35 denaturation cycles at 94°C for 30 seconds, hybridization at 55.7°C for 30 seconds (for ITS1-2 zone amplification) or 58.1°C for 30 seconds (for D1/D2 region amplification), and extension at 72°C for 60 seconds, then a final extension was performed at 72°C for 7 min. The PCR product was verified by performing an 1.5 % agarose gel electrophoresis. The sequencing process was achieved by Macrogen USA, with an automated system based on Sanger’s methodology. The sequences of each amplified zone were analysed using Sequencher 5.4.6 Software (Gen Codes Corporation). Subsequently, Nucleotide BLAST tool (NCBI) was used to obtain the species with the highest homology between the sequences.

### Preparation of the *C. multifidum* broth extract

A culture of *C. multifidium* (1×10^5^ spores/ml) was inoculated into 50 ml of Sabouraud broth (BD-Difco®) and incubated at 30°C and 150 rpm for 2 days. The resulting culture was divided into two parts and transferred to a flask containing 500 ml of dextrose broth and incubated for 3 days at 30°C and 150 rpm. This operation was repeated until obtain 10 L of culture. The broth was separated by vacuum filtration and extracted with ethyl acetate (v/v, 1:1). The organic layers were collected and the solvent removed in a rotavapor resulting in 1.5 g of crude extract.

### Compound isolation

The crude extract (1.5 g) was chromatographed on silica gel by MPLC with a gradient of CH_2_Cl_2_–MeOH (v/v 0:1 to 1:0 v/v) to give 6 fractions (CM1-CM16). Fraction 5 showed the best activity on the bioautography test [19]. It was rechromatographed using silica gel and a gradient of petroleum ether-ethyl acetate (v/v, 90:10 to 80:20) giving as result 8 fractions (CM5.1 – CM5.8), fraction CM5.3 was identify as **2** (6.4 mg), fraction CM 5.5 as **4** (1.7 mg) and fraction CM 5.8 as **6** (8 mg). Fraction CM5.7 was filtered on a Sephadex LH-20 column using CH_2_Cl_2_ as eluent to yield **1** (4.2 mg). Compounds **3** (16.3 mg) and **5** (4.9 mg) were obtained from the purification of fraction CM9 on silica gel using a gradient of petroleum ether-ethyl acetate (v/v, 80:20 a 60:40). A fractionation of CM10 with a solvent system of MeOH-CH_2_Cl_2_ (v/v, 80:20 to 100:0) resulted in the isolation of **7** (5 mg).

### Antibacterial activity assays

The half-maximal inhibitory concentration (IC_50_) was determinate using a microdilution method [20, 21]. Bacteria strains were cultured in 7 ml of Mueller Hinton Broth (MHB) incubated at 37°C for 24 hours. The tested compounds were dissolved in DMSO, final concentration of solvent was less 1%. Dilutions were prepared in 96 well plates mixing prepared DMSO solutions with MHB medium to a final volume of 50 µl. Then, a bacteria culture aliquot of 50 µl was inoculated to the dilutions. Final concentrations of compounds in each well ranged from 500 to 0.98 μg/ml and bacteria density was 5 × 10^5^ CFU/ml. After 24 h of incubation at 37°C the optical density (OD) was read at 595 nm. Tetracycline was used as a positive control in a range of 0.3-0.025 μg/ml. A Probit analysis was performed to determine the IC_50_ of the compounds. The minimum inhibitory concentration (MIC) is determined by observing the first concentration that did not present turbidity or bacterial growth.

## Results and discussions

### Isolation and identification of the active fungus

The culture of solutions prepared from *H. illucens* gut yielded to the isolation of 25 cultivable fungal strains with different morphotypes. The active fungus was identified as *Arthroderma multifidum* with 100% identity and coverage with the AB861747.1 and AB359438.1. This specie is known as the sexual stage (teleomorph) of *Chrysosporium multifidum* (anamorph) [22, 23]. No teleomorph stage was show in the prepared cultures, in contrast it presented abundant pyriform microconidia and hyaline septate hyphae (Fig 1) belonging to its anamorph (*Chrysosporium multifidum*), so we finally named it as *C. multifidum* (GenBank accesion numbers: MK982149 and MK982181). The use of this stage allowed us to determine the antimicrobial activity of its culture supernatant with different experimental methods.

**Fig 1.**
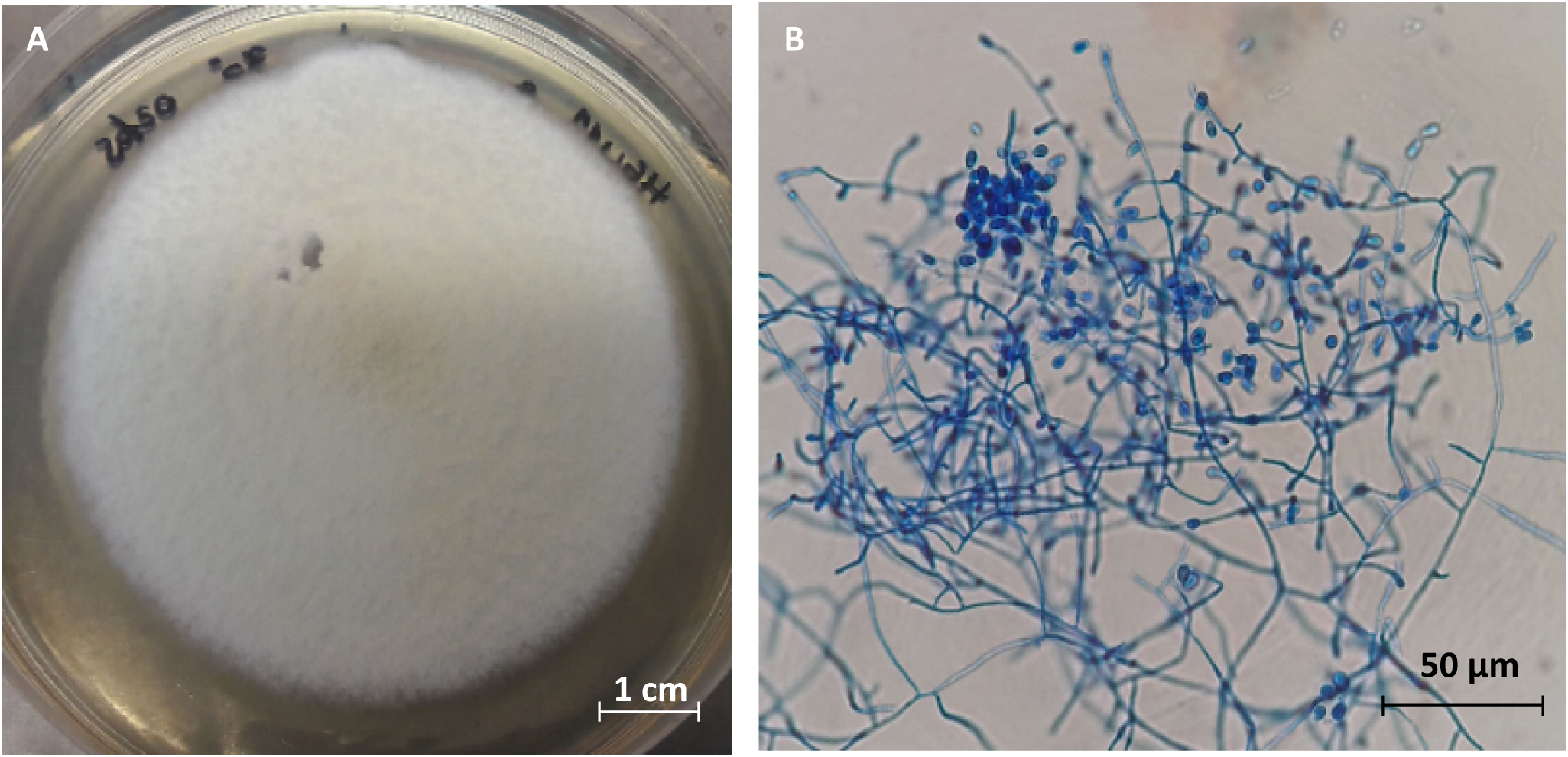
*Chrysosporium multifidum* isolated from *H. illucens* gut after 7 days of incubation at 30°C. Macroscopic view (A). Microscopic view of pyriform microconidia and hyaline septate hyphae (B).

This specie is a saprotroph commonly found in the soil. It has not been studied as an endosymbiont however it is seen as opportunistic and in other cases as a pathogen. Its isolation is directly related to the food consumed, since there are reports that indicate the isolation of the genus *Chrysosporium* from chicken guano samples [24]. This fact increases the probability that they are organisms assimilated along with food (chicken guano diet) and that they would be selected within the fly’s biological system for their convenience (for having enzymes or even beneficial antimicrobial substances) in exchange for providing an environment with enough nutrients for their development [25]. It is known that even some of these selected fungi can survive in glandular cavities or cuticular invaginations called micangias where they can develop and reproduce being favourably transported to new hosts by the insects [26, 27].

### Compound isolation from *C. multifidum* broth extract

The bioguided isolation of ethyl acetate extract prepared from the *C. multifidum* broth resulted in the isolation of: 4-methoxy-2H-pyran-2-one (**1**) [28], 4-methoxy-6-pentyl-2H-pyran-2-one (**2**)[29], 6-(1-hydroxypentyl)-4-methoxy-pyran-2-one (**3**) [29, 30], 6-[(7S,8R)-8-propyloxiran-1-yl]-4-methoxy-pyran-2-one (**4**) [31], pestalotin (**5**) [32, 33], 5,6-dihydro-4-methoxy-6-(pentanoyloxy)-2H-pyran-2-one (**6**) [30, 33] y cyclo-(L-Pro-L-Phe) (**7**) [34]. All the compounds (Fig 2) were identified by comparison of their spectroscopic data (HRMS and ^1^H and ^13^C NMR) with literature, as well as careful examination of their 2D NMR spectra (COSY, HSQC, HMBC). Optical rotations were also coherent with those published, except for (**4**), for which we found an optical rotation close to zero indicating the possible isolation of a racemic mixture (found [α]^20^ D −4.3, c=0.16, MeOH/CH_2_Cl_2_ 9/1; published [α]^25^ D −98.7, c=0.6, MeOH) [31].

**Fig 2.**
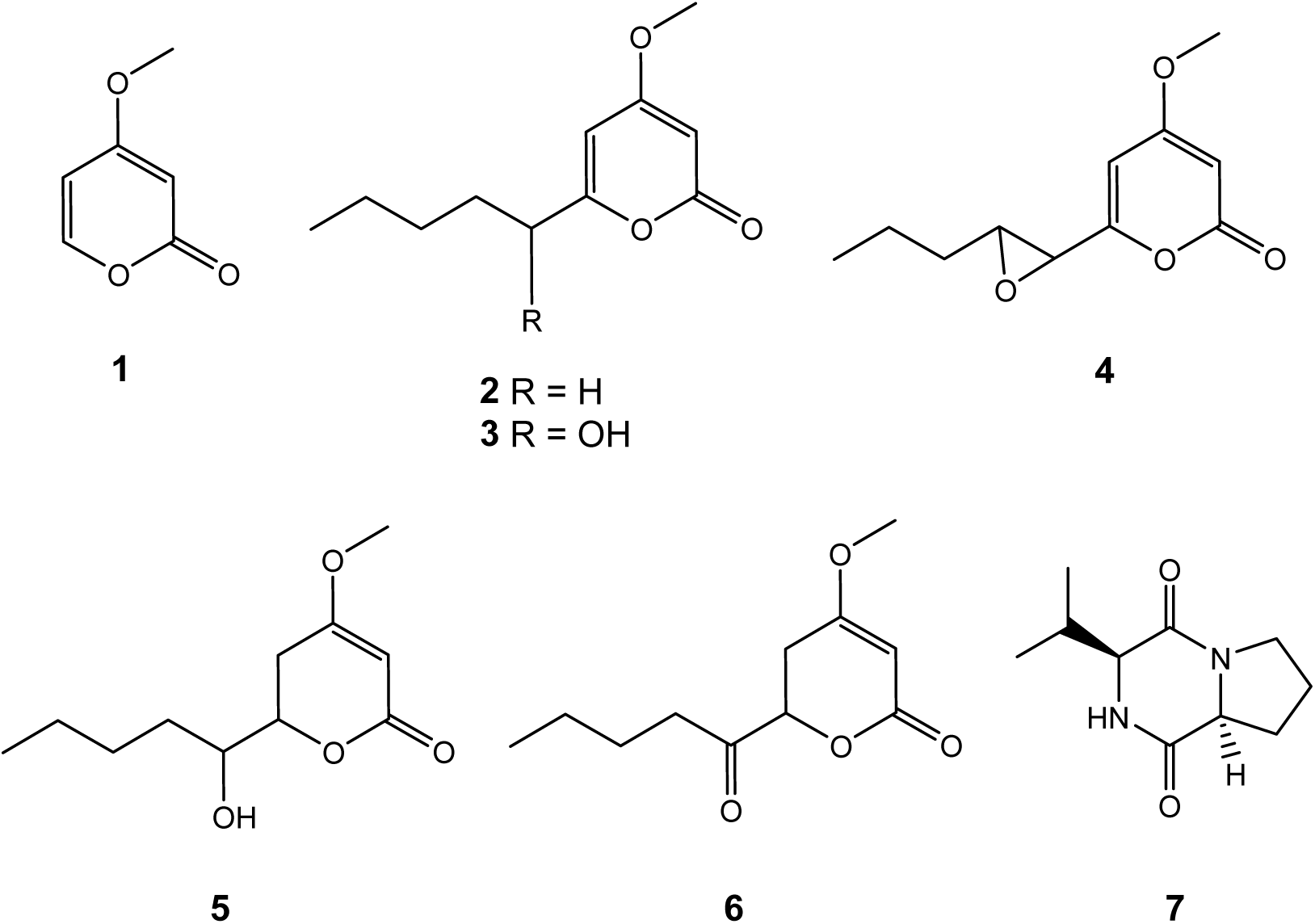
Structure compounds of **1-7** isolated from *C. multifidum* broth extract

This study is the first published chemical characterization of *Chrysosporium multifidum.* The literature describes several chemical prospecting works carried out on species of the *Chrysosporium* genus, which led to the discovery of groups of compounds such as: adenopeptines [35], nucleosides [36], zearalenone derivatives, benzoquinones [38], naphthaquinones [39], anthraquinones [40], benzolactones [41], naphthopyrones [42], naphthalenes [43], phenyl-2(1H)-pyridinones [44], alkylphenols [45], bisdechlorogeodins [46], sterols [47, 48], and caryophillenes [49]. However, there is no prior record of any α-pyrones derivatives, so this would be the first report of these compounds within the genus. Compound **8** is also reported here for the first time within the *Chrysosporium* genus; but this has also been reported from cultures of other fungi and bacteria [50, 51].

### Biological analyses

The α-pyrone **4** showed to be the more active in the bioautography test. Then antimicrobial activity was quantified, results are shown in Table 1. The values of IC_50_ (11.4 ± 0.7 µg/ml) and MIC (62.5 µg/ml) on the methicillin-resistant *Staphylococcus aureus* strain indicate only moderate activity compared to control. The known compounds in *Chrysosporium* genus have displayed biological activites as antitumour [44], antifungal (36, 37, 48, 49) and cytotoxic [41], only naphthaquinon-type compounds isolated from, *queenslandicum* [39] have been shown to be active on gram-positive bacteria *Micrococcus luteus* and *Bacillus subtilis* with MIC values close to those obtained in this work. On the other hand, both natural and synthetic α-pyrones have shown antimicrobial and antifungal activity on a variety of species [52, 53]. Substitutions in positions 4 and 6 of the pyrone ring would be related to this activity. Subsequent trials should be carried out to test the activity of all derivatives of isolated α-pyrones on other groups of bacteria including gram-negative ones.

**Table 1.** Antimicrobial activity of compound **4** against MRSA strain

| Compound | IC<sub>50</sub> (μg/mL) | MIC (μg/mL) |
| --- | --- | --- |
| 4 | 11.4 ± 0.7 | 62.5 |
| Tetracycline | 0.1 ± 0.02 | 0.4 |

## Conclusion

A total of 25 fungi colonies were isolated from the gut of *Hermetia illucens* larvae fed with chicken guano. These colonies were tested on methicillin-resistant *Staphylococcus aureus* (MRSA) ATCC 43300 and *Salmonella* Typhimurium ATCC 13311. One colony showed the best activity on the MRSA strain, this specimen was subsequently identified as *Chrysosporium multifidum*. A broth culture of the fungus was prepared and seven compounds were isolated using chromatographic methods and bioguidage by bioautography. The active compound against MRSA was identified as the α-pyrona **4** with a MIC of 62.5 µg/ml. These first results in the exploration of the microbiota of *H. illucens* open a path to understand the interaction of this fungus with other microorganisms that allow the larva to control the pathogenic microbes introduced through its food made of contaminated organic waste.

## Author Contributions

### Conceptualization

Michel Sauvain, Yesenia Correa, Denis Castillo.

### Investigation

Yesenia Correa, Billy Cabanillas, Valérie Jullian, Daniela Álvarez, Denis Castillo, Cédric Dufloer, Beatriz Bustamante,, Michel Sauvain.

### Funding acquisition

Michel Sauvain.

### Methodology

Elisa Roncal, Edgar Neyra, Patricia Sheen, Cédric Dufloer

### Supervision

Billy Cabanillas, Michel Sauvain, Yesenia Correa, Valérie Jullian.

### Validation

Billy Cabanillas, Michel Sauvain, Yesenia Correa, Valérie Jullian.

### Writing - original draft

Billy Cabanillas, Yessenia Correa, Michel Sauvain, Daniela Alvarez.

### Writing - review & editing

Billy Cabanillas, Yessenia Correa, Michel Sauvain, Daniela Alvarez

